# Reaching for known unknowns: Rapid reach decisions accurately reflect the future state of dynamic probabilistic information

**DOI:** 10.1101/2020.07.31.231563

**Authors:** Nathan J. Wispinski, Scott A. Stone, Jennifer K. Bertrand, Alexandra A. Ouellette Zuk, Ewen B. Lavoie, Jason P. Gallivan, Craig S. Chapman

**Affiliations:** Department of Psychology, University of Alberta, Edmonton, Alberta, Canada; Faculty of Kinesiology, Sport, and Recreation, University of Alberta, Edmonton, Alberta, Canada; Neuroscience and Mental Health Institute, University of Alberta, Edmonton, Alberta, Canada; Centre for Neuroscience Studies, Queen’s University, Kingston, Ontario, Canada; Department of Psychology, Queen’s University, Kingston, Ontario, Canada; Department of Biomedical and Molecular Sciences, Queen’s University, Kingston, Ontario, Canada

**Keywords:** Motor control, Prediction, Reaching, Probability, Visuospatial attention

## Abstract

Everyday tasks such as catching a ball appear effortless, but in fact require complex interactions and tight temporal coordination between the brain’s visual and motor systems. What makes such interceptive actions particularly impressive is the capacity of the brain to account for temporal delays in the central nervous system—a limitation that can be mitigated by making predictions about the environment as well as one’s own actions. Here, we wanted to assess how well human participants can plan an upcoming movement based on a dynamic, predictable stimulus that is not the target of action. A central stationary or rotating stimulus determined the probability that each of two potential targets would be the eventual target of a rapid reach-to-touch movement. We examined the extent to which reach movement trajectories convey internal predictions about the future state of dynamic probabilistic information conveyed by the rotating stimulus. We show that movement trajectories reflect the target probabilities determined at movement onset, suggesting that humans rapidly and accurately integrate visuospatial predictions and estimates of their own reaction times to effectively guide action.

## 1. Introduction

Humans exist in a dynamic world. Everyday tasks such as walking onto a moving escalator or catching a ball appear simple, but require tight temporally-coupled communication between visual and motor areas of the brain to ensure the action is successful. A key aspect of both of these tasks is that they require interception—demanding that the person get their body to the right place at the right time. To have this kind of successful interaction with our environment, predictions about the future state of moving objects must be computed by the brain and transformed into action. Catching a ball, for example, requires that the visual representation of the ball and its likely trajectory be transformed into the appropriate arm and hand movements, ultimately producing an anticipatory interceptive movement based on predictive internal models of object acceleration and gravity (1–3).

What makes such interceptive actions particularly impressive is the capacity of the brain to account for the various temporal limitations of the central nervous system. Visuomotor processes, involving sensory evidence integration, action planning, and movement initiation, are subject to neurophysiological transmission delays ranging from 100 to 450 ms (4–6). Given our natural aptitude to intercept moving objects (1, 7–10) even when their motion can’t be fully observed (9, 11, 12), theories articulate that we must pre-plan (6, 13), adjust on the fly (14), or mix planning and adjustment (15) to overcome these delays and produce successful interception actions. Empirically, we see evidence that the brain predicts the delays of sensory inputs in visual illusions. For example, in the flash-lag effect (16), a predictably moving object is perceived as occupying its future location. Likewise, we see evidence that the brain predicts the delays of motor outputs during decision making. For example, during a random dot motion task, neuronal activity thought to reflect the decision variable terminates ∼50 ms before movement initiation (17).

Studies of interception tasks have shown that humans are adept at predicting the future location of an object based on the movement of *that* object (e.g., by continuing smooth pursuit of an object through a period of occlusion (9), fixating on the object intended for interception (7, 8), or hitting a ball with a bat (1)). Yet, anecdotally, we also know that humans can make predictions about where to move based on other objects in the environment (e.g., obstacles) (18, 19), and plan actions toward locations where the eyes are *not* fixated (e.g., anti-pointing tasks) (20–22). An intuitive example is a hockey forward who shoots opposite the position of the goalie to score a goal. Here, the already complex sensory-to-motor transformation must introduce yet another mediating cognitive variable—the representation of where the goalie *will not be* based on where the goalie will be. Here, we wanted to assess this particular capacity—how well can participants plan an upcoming movement based on a dynamic, predictable stimulus that is *not* the target of action.

One tool for assessing dynamic cognitive states is to analyze the shape of movement trajectories (4, 23–31). Hand or computer mouse trajectories can reflect the deliberation of external information, such as random dot motion stimuli (4, 5), number magnitude (32, 33), or word processing (28). Fluctuations in movements toward a final choice can also reflect internal information, such as the subjective value of snack foods (34). Typically, movement trajectories that curve between potential targets suggest conflict or indecision, while trajectories relatively straight toward a target reflect less competition between alternatives (27, 31, 35).

Of particular note, movements can reflect static or changing probabilities of multiple potential targets in space. When required to reach toward one of many potential targets on a screen, movement trajectories are sensitive to target number, suggesting a rapid integration of static probabilistic information during movement planning (23, 36, 37). This information can bias movement trajectories even when movements need to be initiated less than 325 ms after stimuli onset (36). Others have shown that movement trajectory planning can also incorporate changing probabilistic information over time (4). In one group of studies, subjects were asked to reach toward a left or right target to indicate whether a group of dots on a screen are moving left or right. In this task, dot motion is noisy, so motion information fluctuates from timepoint-to-timepoint. In these studies, initial movement trajectories reflect fluctuating dot motion information that occurred roughly 350 ms in the past (4, 5).

Here we questioned the extent to which movement trajectories also convey internal predictions about the future state of dynamic probabilistic information. To examine this, we manipulated probabilistic information dynamically between two potential targets. Participants were presented with a stimulus that rotated in a circle and were required to launch a movement towards the potential targets prior to the final target being cued (i.e., go-before-you-know task). Critically, the position of the stimulus at movement onset determined how likely each of the two potential targets would be selected as the ultimate target of action on that trial. Previous work has shown that central (endogenous) versus peripheral (exogenous) cue-stimuli elicit different patterns of prediction and evolve over different time courses (38). To examine if this affected our dynamic prediction task, we collected data from two groups of participants—one where the rotation stimulus was an arrow that rotated about the central fixation and one where the rotation stimulus was a box that moved on a more peripheral path adjacent to the potential targets. We show that, across both groups, movement trajectories reflect target probabilistic information determined at movement onset, suggesting that humans rapidly and accurately integrate visuospatial predictions and estimates of their own reaction times to effectively guide action.

## 2. Materials and methods

### 2.1. Overview of procedure

Humans use predictions to overcome sensorimotor delays such that successful actions are generated to intercept moving objects. Here we use an analysis of behavior (accuracy and movement trajectories) in a rapid reach task to test whether these same predictive capacities extend to movement planning based on predictable, but dynamic sensory evidence. The stimulus in our experiments is separate from the target and conveys information about the probability of the final target location, rather than cueing location directly. In this task, we extend our previous go-before-you-know paradigm (23, 39–42) which requires participants to initiate a movement in response to a go-signal before one of two potential target locations is revealed as the final target. Here, the probability of the upcoming target location was conveyed to participants via a stimulus that rotated at a fixed rate, either clockwise (CW), counter-clockwise (CCW), or, on baseline/control conditions, remained stationary (Fig. 1; see videos linked in Data and code availability). To test for possible differences in endogenous versus exogenous cueing (38), the stimuli conveying probability used in the present study differed across two groups: one group saw a central red arrow, and the other group saw a more peripheral red box, both of which rotated around the central fixation cross (Fig. 1b). The position of the probability-stimulus at movement onset dictated the probability with which one of two targets was selected as the final target for action (Fig. 1d). Thus, have the highest chance for success participants needed to be monitoring and predicting from the rotating probability stimulus before the go-signal *and* during reaction time.

**Fig. 1.**
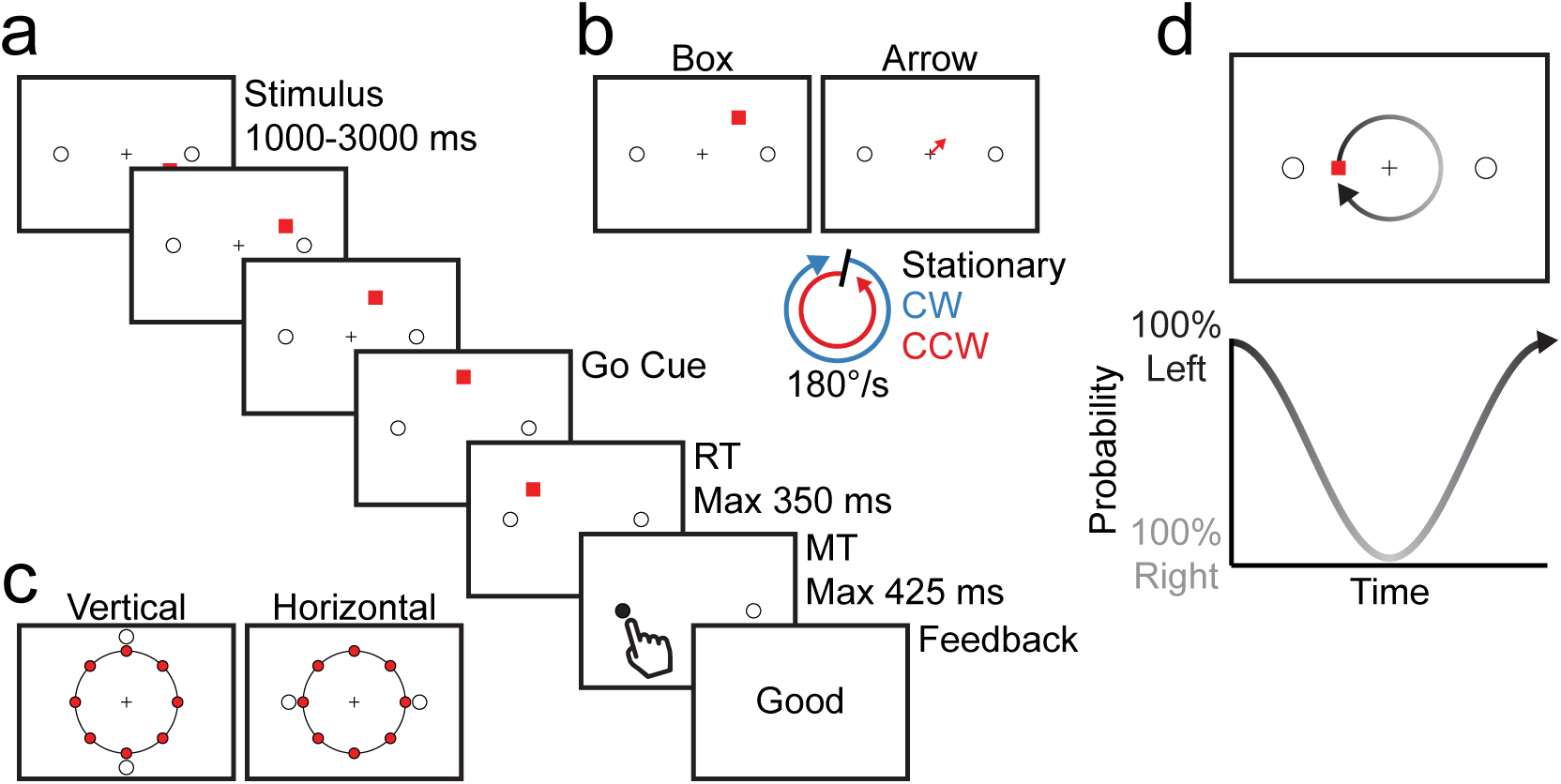
Stimuli and trial sequence. Also see videos linked in Data and code availability. (a) Example trial where a stimulus rotates counterclockwise (CCW) around the fixation. Once the stimulus hits a predetermined point along the circle (e.g., at the very top of the circle), the fixation disappears and a ‘beep’ plays. The stimulus continues rotating until the participant lifts their finger off the start button, after which the stimulus disappears and the final target is cued. Participants must begin their rapid reach before knowing which of the two targets will be cued. (b) The stimulus that determined target probability was a box or an arrow for different sets of participants. (c) Targets were arranged vertically or horizontally on each trial. One of eight equally-spaced points along the circle was pre-determined for each trial at which the go-signal would occur (depicted here as red circles). (d) As the stimulus moves (top panel, shown clockwise), the probability of target location oscillates (closer to left, black, left target; or closer to right, grey, right target).

Since the stimulus conveying target probability moves in a circle, the probability of any one target being selected varies sinusoidally (Fig. 1d). We capitalized on this sinusoidal feature of target probability to test for sinusoidal characteristics of behaviour in our key dependent measures—choice accuracy and reach curvature (indexed by reach area; Fig. 2). The trials in which the probability stimulus remained stationary serve as the starting point for our analysis (black curves in Fig. 3). In these stationary trials, we predict and find that participants are most accurate and reach trajectories are most straight (low area, e.g., grey trajectory in Fig. 2b) when the probability-stimulus perfectly predicts the target location (i.e., 100% probability), and least accurate and least straight (high area, e.g., green trajectory in Fig. 2b) when the probability stimulus is ambiguous with respect to final target location (i.e., 50% probability). To test for participants’ ability to use the rotating probability-stimulus to guide action planning we compare the rotating trials (CW in blue, CCW in red; Fig. 3) to these stationary trials.

**Fig. 2.**
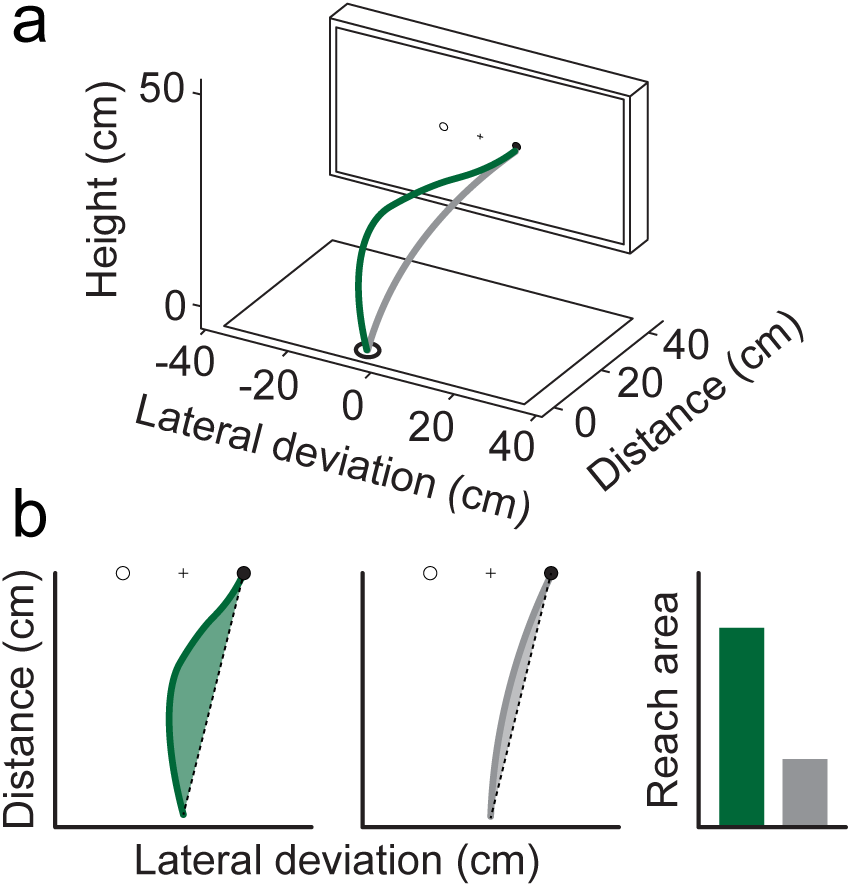
(a) Example of the three-dimensional reach trajectories collected. (b) Examples of two reach trajectories on trials where the right target is cued. The area between the reach trajectory, and a straight line from the start position to this participant’s mean endpoint for right target trials, is used to index reach curvature. When reach trajectories travel between the two targets the reach area is larger (green), and when trajectories are travel straight to one target the area is smaller (grey).

**Fig. 3.**
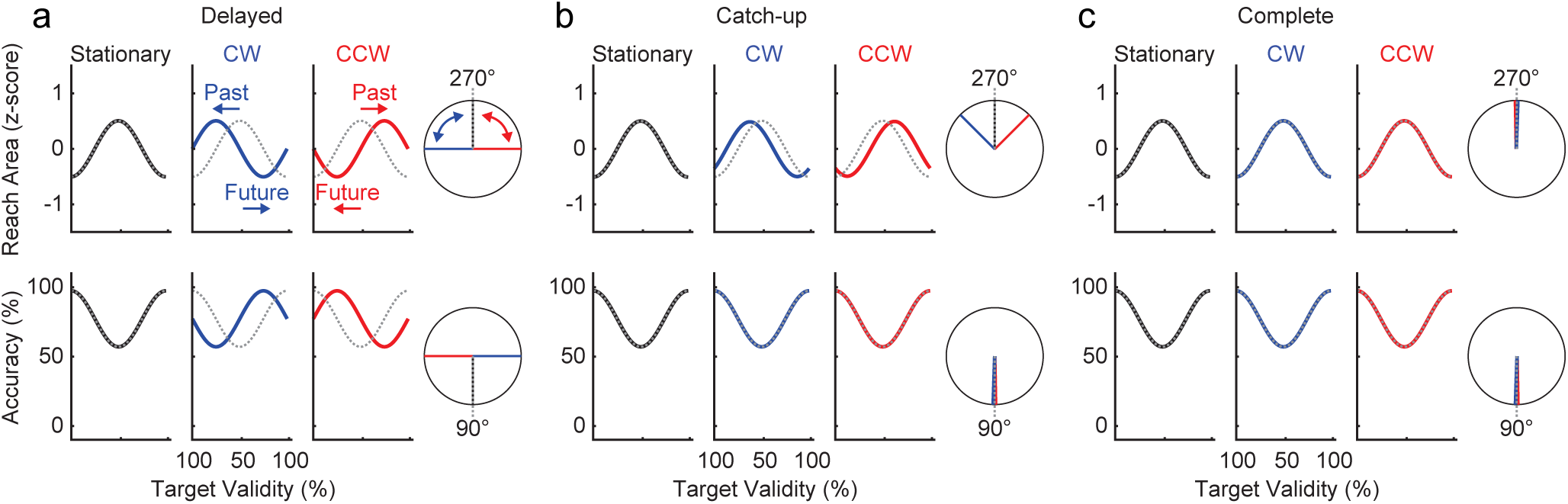
Predictions from left to right. Sinusoids for stationary (black), clockwise (blue), and counterclockwise (red) trials show predictions for reach area (top) and accuracy (bottom). Circles next to sinusoid panels are polar plots, with condition-colored lines showing the predicted phase offsets relative to an expected phase (grey dotted lines). (a) Delayed. No predictive processing, or insufficient prediction. Accuracy and reach area reflect target probability in the past. (b) Catch-up. Reach area reflects target probability in the past, but information is used during the movement to correct the reach so that accuracy reflects the final target probabilities. (c) Complete. Prediction is accurate and fast. Information about target probability is able to be used at the time of movement onset, when target probability is actually determined, and is not biased by other factors. Reach Area and Accuracy both reflect the final target probability.

Our results test between three patterns of hypothesized behaviour (Fig. 3). First, as a baseline, we show what we would expect to see if participants were not predicting the future location of the rotating stimulus, but rather, “living in the past” (Delayed, Fig. 3a). Here, the CW and CCW data would show a shifted sinusoid whose phase (polar plots; Fig. 3) is out of alignment (specifically, delayed in time) for both accuracy and reach area. In this case, accuracy and reach area would reflect a temporally-outdated location of the probability-stimulus. Under this prediction, behavioural measures could reflect target probability determined at a salient event like the go-signal, or at a constant delay reflecting computation and transmission delays. This result would be consistent with data from unpredictable stimuli like in a random dot motion task, where movement trajectories and accuracy reflect the status of a decision variable several hundred milliseconds in the past (4).

Second, if we imagine that participants are living in the past at the onset of movement but use the time available during the executed movement to make online corrections (e.g., changes of mind) (4), we would predict a “catch-up” pattern of results (Fig. 3b). Here, the phase of the sinusoid for the reach area of CW and CCW trials lags the phase of the reach area across static trials, but the phase of the sinusoid for accuracy “catches up” such that all across-trial phases align. In this case, participants initially aim toward an outdated probabilistic location, but successfully correct their movements in flight to reach and touch the final target.

Finally, third, if we imagine that participants are successfully predicting the future probability at the moment of movement onset (and thus, accurately accounting for sensorimotor processing delays and being unbiased by other factors), we would expect to observe a “complete” pattern of results (Fig. 3c). Here the CW and CCW data would match the stationary data. That is, even though rotation trials are dynamic, the prediction is accurate, rapid, and updated in real-time such that participants both aim toward an up-to-date probabilistic location and correctly touch the final target.

While the measures of reach behaviour described above are the focus of this study, we can also test how the sinusoidal nature of target probability might induce sinusoidal changes in reaction time. In tasks requiring action in response to targets of varying uncertainty, participants have been shown to adjust movement and reaction times to improve visuomotor accuracy in trials with greater uncertainty (43). We would therefore predict reaction times to fluctuate sinusoidally with target probability. Specifically, when anchored to the go-signal, we would predict trials where target uncertainty is high (probability ≈ 50%) to result in longer reaction times, with participants maximizing the amount of visual evidence accumulated in support of final target probability. In contrast, we would predict trials where the target uncertainty is low (probability ≈ 100%) to result in shorter reaction times, potentially allowing participants to decrease motor errors by increasing movement time.

### 2.2. Participants

Twenty-seven participants (19 women; Age: *M* = 22.78, SD = 4.19) took part in the arrow experiment, while twenty-eight participants (13 women; Age: *M* = 22.96, SD = 3.53) took part in the box experiment. All participants provided written consent before the experiment, and were compensated with course credit for participation. Experimental procedures were approved by Western University’s Research Ethics Board. Only data from right-handed participants with normal or corrected-to-normal vision were analyzed.

### 2.3. Equipment and stimuli

Participants sat in front of a 40” touchscreen (NEC MultiSync^©^ LCD4020 refresh rate 60 Hz; Fig. 2a), and made rapid reaching movements to targets on the screen (see videos linked in Data and code availability). Two active infrared markers were taped to the participant’s right index finger, and tracked reaching movements throughout the experiment (Optotrak, 150 Hz). All stimuli presentation and data collection were controlled with MATLAB (The Mathworks, Natick, MA) using Psychtoolbox (Version 3) (44–46).

### 2.4. Trial sequence and procedure

Participants performed a variant of a go-before-you-know task (23, 47), requiring them to initiate a rapid reach movement before they knew which of two potential targets would be cued as the final target. The current study involved the presentation of a box (box experiment) or arrow (arrow experiment) stimulus that could either rotate around a central fixation (clockwise or counterclockwise) or remain in a fixed position (stationary). Two potential targets were presented (placed horizontally or vertically), and after a variable delay, an auditory beep would signal the participant to begin their reaching movement. At movement onset, one of the two targets was cued as the final target—the probability of which was determined by the location of the probability-stimulus at movement onset (Fig. 1d). Participants were informed that the final location of the stimulus dictated target probability prior to commencing the task, and were given practice trials until they reported feeling comfortable with the experimental procedure (e.g., timing constraints).

Trials began with the participant holding down the start button (Fig. 2a, positioned 5 cm from the front edge of the table) with their right index finger. The start button was placed so that participants would need to reach forward 40 cm and up 25 cm to touch the center of the screen in front of them.

With the start button held down, a central fixation cross would appear with two targets on a screen with a white background (Fig. 1a). The targets on each trial were arranged either horizontally or vertically, evenly counterbalanced across all trials (Fig. 1c). Potential targets were black outlines of circles 2 cm in diameter, and located 9 cm from the fixation cross at the center of the screen. Participants were instructed to maintain central fixation at all times during the experiment.

Next, a stimulus would appear. For participants in the box experiment, this stimulus was a red square 2 cm wide (Fig. 1b). For participants in the arrow experiment, this stimulus was a red arrow with its base at the fixation, and extending ∼2.2 cm outward. On stationary trials, the stimulus would appear located at, or pointing toward, one of 8 evenly-spaced locations 7 cm from the origin (0°, 45°, 90°, 135°, etc.; evenly counterbalanced across trials, Fig. 1c) and not move throughout the trial. On non-stationary trials, this stimulus would appear at, or point toward, one of 120 evenly-spaced points centered 7 cm from the origin, with the start location of the stimulus chosen from a random uniform distribution. During these trials, the stimulus would rotate either clockwise or counterclockwise about the fixation along (box experiment), or pointing toward (arrow experiment), an invisible circle with 7 cm radius at a constant angular velocity of 180°/s. For both trial types (stationary and non-stationary), the stimulus remained on the display until participants initiated their reaching movements in response to a go-signal. The go-signal consisted of an auditory beep paired with the simultaneous disappearance of the central fixation cross.

On stationary trials, this go-signal always occurred one second after the onset of the stimulus. On non-stationary trials, the stimulus would rotate around the origin for a minimum of one second, but would continue moving until it had reached one of the eight predetermined locations on the circle (0°, 45°, 90°, 135°, etc.; evenly counterbalanced across trials; Fig. 1c). Once the box or arrow stimulus had reached this specified location, the participant would be signalled (via fixation disappearance and the coincident beep) to initiate their movement.

Participants had 350 ms (box experiment) or 325 ms (arrow experiment) after the go-signal to lift their finger off the start button. Upon successful button release, the stimulus disappeared and one of the two circles was filled in. Participants then had 425 ms to touch the cued final target on the screen (i.e., we required that the reach movement be ballistic). The probability of a target filling in was based on the location of the stimulus when the start button was released.

For example, a target had a 100% probability of being cued as the final target if the probability-stimulus was located directly next to it (box) or pointed directly towards it (arrow) at reach onset. If the probability-stimulus was halfway between the targets when the reach was initiated, both targets had a 50% chance of being filled in (Fig. 1d). At the end of each trial, participants received feedback on their performance. If participants lifted their finger earlier than 100 ms after the go-signal (i.e., the reach movement was anticipatory), a “Too Early” error message would be presented after trial completion. If participants exceeded the reaction time limit, or the 425 ms movement time limit, the trial would similarly end with a “Time Out” or “Too Slow” error message, respectively. Participant accuracy was denoted by either a “Miss” message should they have touched the screen outside of a 6 cm x 6 cm invisible box centered on the correct final target, or a “Good” message should they successfully complete the trial without any errors.

Trials were equally counterbalanced for target arrangements (horizontal or vertical), stimulus motion (stationary, clockwise, or counterclockwise), and stimulus position at the time of the go-signal (eight equally-spaced positions around the origin; Fig. 1c). As such, there were 48 unique conditions (2 target arrangements x 8 trigger positions x 3 rotations), each repeated 12 times for a total of 576 trials. Trial order was fully randomized for each participant.

### 2.5. Pre-processing

Trials were deemed as usable for analysis if they were not “Too Early” or “Time Out” trials, did not contain movement recording errors, or did not contain “out of bounds” start or end positions. Additionally, participants were rejected for analysis if they had 25% or fewer usable trials in 8 or more of the 48 unique conditions. These rejected participants are not discussed further. This criterion was enforced so that participants had at least three trials in most conditions for analysis. Three subjects were rejected from the box experiment, while six subjects were rejected from the arrow experiment. One subject was also rejected from each experiment for initially reaching backward off the start position in the majority of trials, leaving n = 24 and n = 20 for each the box and arrow experiments, respectively.

Data cleaning and trial rejection were conducted following the recommendations in Gallivan & Chapman (2014) for rapid reaching experiments. In brief, reach trajectories were space-normalized to 200 equally-spaced points along the ∼40 cm distance from the start position to the screen (47). Reach area was calculated as the approximate area between a reach trajectory on a correct trial and the straight line between the start position and average endpoint for that corresponding target (left, right, up, down) calculated for each subject (Fig. 2b; for previous use see (48, 49). Area was calculated in two-dimensional space along the axis of interest on that trial (e.g., horizontal axis for horizontal target trials). Reach areas were then z-scored for each subject within each target orientation condition (left, right, up, down). Reach area normalization was performed because biomechanical differences within and between subjects created differences in reach area for different reach directions that were not of interest in this study. Larger normalized reach areas correspond to trajectories that move more in between the two targets, whereas smaller reach areas correspond to trajectories that more closely follow a straight line path to the correct, filled-in target. As such, reach area can be used to estimate the level of competition or indecision between several potential targets in space (31, 47, 50).

Reaction time was calculated as the time from the go-signal auditory beep to the release of the start button. Movement time was calculated as the elapsed time between button release and when a touch was detected on the touchscreen. Unlike “Too Early”, “Time Out”, and “Miss” trials, we did not automatically reject “Too Slow” trials. Instead, “Too Slow” trials >2 SD above a participant’s mean (after excluding all trials with a movement time >850 ms) were rejected for analysis.

Errors on each trial could be a combination of “Too Early” (*M* = 1.02%, Range: 0% - 5.21%), “Miss” (*M* = 8.08%, Range: 0.52% - 20.31%), “Time Out” (*M* = 12.02%, Range: 0.35% - 32.12%), >2 SD of mean movement time (*M* = 4.06%, Range: 1.22% - 7.64%), reaches with recording errors (*M* = 1.21%, Range: 0% - 9.72%), and reaches with “out of bounds” start or end positions (*M* = 6.49%, Range: 0.17% - 32.29%). In total, participants whose data was analyzed had a mean of 86.02% usable trials for analysis (Range: 57.81% - 98.96%), and of those trials a mean of 86.06% were correct (Range: 72.59% - 96.97%). These trial rejection numbers are generally in line with recommendations for rapid reach experiments (47).

### 2.6. Model

To overcome sparsity of sampling (there were 120 possible stimulus locations) and to directly test for the predicted sinusoidal patterns of data (Fig. 3), we reduced the data collected in this experiment by fitting a sine wave model to each condition for each subject. The sine wave model consisted of a fixed period equal to the rate of stimulus rotation (180°/s), and three free parameters: mean shift (*µ*), amplitude (*A*), and phase shift (*ϕ*).

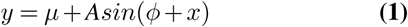

To fit data to this sine wave model, circle positions on vertical target trials were rotated 90° so that they would line up with horizontal target trials (i.e., 100% target probability occurred at the same circle location for horizontal and vertical trials). Circle positions were then collapsed so that positions started at 100% target probability of left targets, decreased to 50% target probability, and then ended at 100% probability for right targets (Fig. 1d). For each subject and for each condition (e.g., Fig. 4a shows a subject in the box experiment, horizontal targets, and clockwise stimulus rotation), single trial data were fit to the sine wave model using a least squares cost function. One-hundred fits were performed using the fminsearchbnd function in MATLAB with random initial parameters, and the fit with the lowest cost was taken as the final parameter estimate. The amplitude parameter was constrained to be higher than zero for all fits, as it caused inaccurate phase parameter estimates if amplitude was too low.

**Fig. 4.**
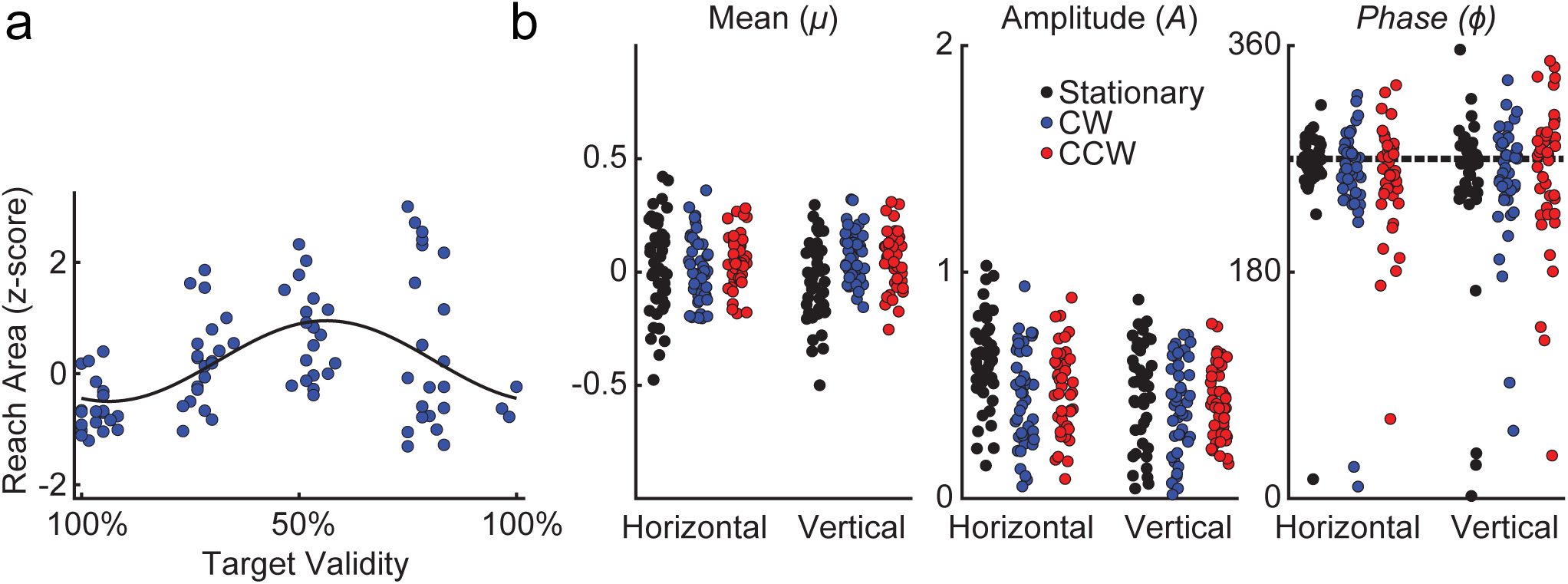
Fitting sine wave models to reduce data. (a) Example of a single subject, single condition sine-wave fit where each data point is a single trial. Sine waves with fixed period (matching the probability profile of the rotating stimulus), variable mean shift, amplitude, and phase shift were fit to single-trial data. Shown is normalized reach area by target validity locations for a single participant in the box experiment when targets were arranged horizontally and the probability stimulus was rotating clockwise, *R*^2^ = 0.23. (b) Sine wave parameter fits to normalized reach area where each data point is one subject’s data for each parameter (*µ* - left panel, *A* - middle panel, *ϕ* - right panel) in each condition (Stationary - black, CW - blue, CCW - red) and experiment (Box and Arrows). Dashed line in the phase panel represents the expected phase for normalized reach area (lowest at 100% target validity, highest at 50% target validity).

By fitting sine waves to each condition for each participant, these data were reduced to three parameters (mean, amplitude, and phase), which were used for statistical comparisons (Fig. 4b). Overall, these sine waves are reasonable descriptors of the data and provided useful data reduction. First, the model period corresponds directly to the independent variables of stimulus motion and changes in target probability with location (i.e., 180°/s). Second, the fitted models describe the dependent variables in different target probability locations reasonably well, given that the sine wave model is fit to single-trial data (reaction time, mean *R*^2^ = 0.09, range: -0.26 - 0.45; accuracy, mean *R*^2^ = 0.08, range: 0 - 0.37; reach area, mean *R*^2^ = 0.13, range: -0.01 - 0.45). Reach area and reaction time were only calculated for correct trials.

### 2.7. Statistical analysis

#### 2.7.1. Phase

Our primary theoretical motivation was to test whether prediction of probability would be evident in our dependent measures. As such, of our model-fitted dependent measures, the phase parameter is of the most theoretical importance. However, estimated phase parameters reasonably match a circular normal distribution, which violates assumptions of many statistical tests, such as a linear repeated-measures ANOVA. Therefore, the phase parameters for the sine waves fit to each of reaction time, reach area, and accuracy were compared using circular statistics (51). In particular, we were interested if estimated phases in each condition were significantly different from an expected phase. For instance, in the stationary stimulus condition, we would expect reaction times to be the fastest, reach area to be the smallest (reaches most straight), and accuracy to be highest when target probability was 100%. We expect the reverse pattern when the probability was 50% (slow reaction times, large reach areas, and low accuracy). Below we compare whether the observed phase estimates in each condition were significantly different from the expected phase using one-sample circular t-tests. In addition, we wanted to know how each of our stimulus conditions differed from one another. So, we also ran all possible circular paired t-tests of stationary vs. clockwise vs. counterclockwise stimulus conditions. This led to 18 total circular t-tests (3 dependent measures x (3 one-sample + 3 paired)), which were Bonferroni-corrected to a statistical threshold of 0.0028 (i.e., 0.05/18). Our investigation of phase collapses across the other factors in our experiment (Experiment: Box or Arrow, and Target Arrangement: Vertical or Horizontal) because our main theoretical questions are driven by Rotation.

#### 2.7.2. Mean and Amplitude

For mean and amplitude parameters estimated from each dependent variable (reaction time, reach area, accuracy), we conducted a 2 (target arrangement: horizontal vs. vertical) x 3 (rotation: stationary vs. clockwise vs. counterclockwise) x 2 (experiment: box vs. arrow) repeated-measures ANOVA. All main effects and interactions were Greenhouse-Geisser corrected, and also corrected using a sequential Bonferroni procedure to control for the familywise error rate of all the repeated-measures ANOVA tests together (52).

## 3. Results

Accuracy and reach area were analyzed relative to the final target probabilities on each trial (i.e., when the probability-stimulus disappeared at the beginning of a movement). However, reaction time was analyzed relative to the target probabilities at the go beep. Locking reaction times to the go beep can give us a picture of how target probabilities influence movement onset times.

### 3.1. Effects of rotation on phase

As articulated in our Methods, our primary motivation was to analyze the effect of rotation condition on the estimated phase parameters of the data. These analyses speak to whether the sinusoidal pattern of the dependent measures are shifted depending on whether the stimulus was stationary or rotating, and should indicate whether reach behaviour reflects a delayed, catch-up, or complete sensorimotor prediction process based on dynamic target probability (Fig. 3).

For reaction time data (Fig. 5), we find that the distribution of estimated phases when the stimulus is stationary is not different from the expected phase (phase difference = 2.65°, *p* > 0.0028). Here we test against an expected phase where fastest reaction times occur when probability is 100% and slowest reaction times occur when probability is 50%. However, estimated phases in conditions where the targets are moving clockwise (phase difference = 51.16°) or counterclockwise (phase difference = -63.94°) are significantly different from the expected phase (*p*s < 0.0028). Pairwise comparisons indicate that both the stationary and clockwise phases are significantly different from the counterclockwise phase (*p*s < 0.0028). However, stationary and clockwise phases are not significantly different from each other (Fig. 5; *p* > 0.0028). This pattern of results suggests that when the probability-stimulus is rotating, participants are reacting to the probability state that the stimulus is *approaching*, rather than reacting to where the probability-stimulus is actually located at the time of the go-signal.

**Fig. 5.**
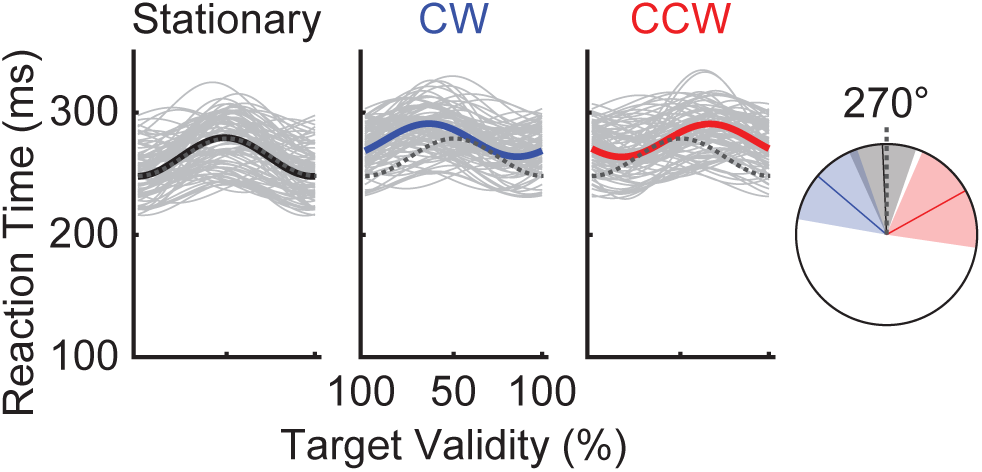
Sine wave models fit to reaction time data over target validity positions, time-locked to the go beep. Sine waves from each individual condition (e.g., a subject in the box experiment, horizontal targets, clockwise rotation) are in light grey and a sine wave with average mean, amplitude, and phase parameters are in solid colors (black for stationary, blue for clockwise, red for counterclockwise). On the right is a circle showing a polar plot describing the phase parameters and their confidence intervals (Bonferroni-corrected 95%). The sinusoidal pattern of results corresponding to the expected phase are plotted dashed grey lines. Average stationary phase was not significantly different from the expected phase, while phases in the rotating conditions were significantly different from the expected phase.

For reach area data (Fig. 6), the expected phase is that reach area would be smallest when probability was high, and largest when probability was low. All estimated reach area phases do not differ from the expected phase regardless of if the stimulus was stationary (phase difference = 2.39°), moving clockwise (phase difference = -5.61°), or moving counterclockwise (phase difference = -5.11°, all *p*s > 0.0028). For accuracy data (Fig. 6), the expected phase is that accuracy would be highest when probability was high, and lowest when probability was low. All estimated accuracy phases do not differ from the expected phase regardless of if the stimulus was stationary (phase difference = -8.85°), moving clockwise (phase difference = 3.91°), or moving counterclockwise (phase difference = -0.15°, all *p*s > 0.0028). Pairwise comparisons indicate that stationary, clockwise, and counterclockwise phases are not significantly different from each other (*p*s > 0.0028) for both reach area and accuracy data (Fig. 6). This pattern of results shows that people were accounting for sensorimotor delays and building those sensorimotor delays into their reach planning. This aligns with our Complete prediction hypothesis (Fig. 3) and demonstrates that, in this task, predictive mechanisms were being successfully deployed based on a probability-stimulus that was separate from the actual target location.

**Fig. 6.**
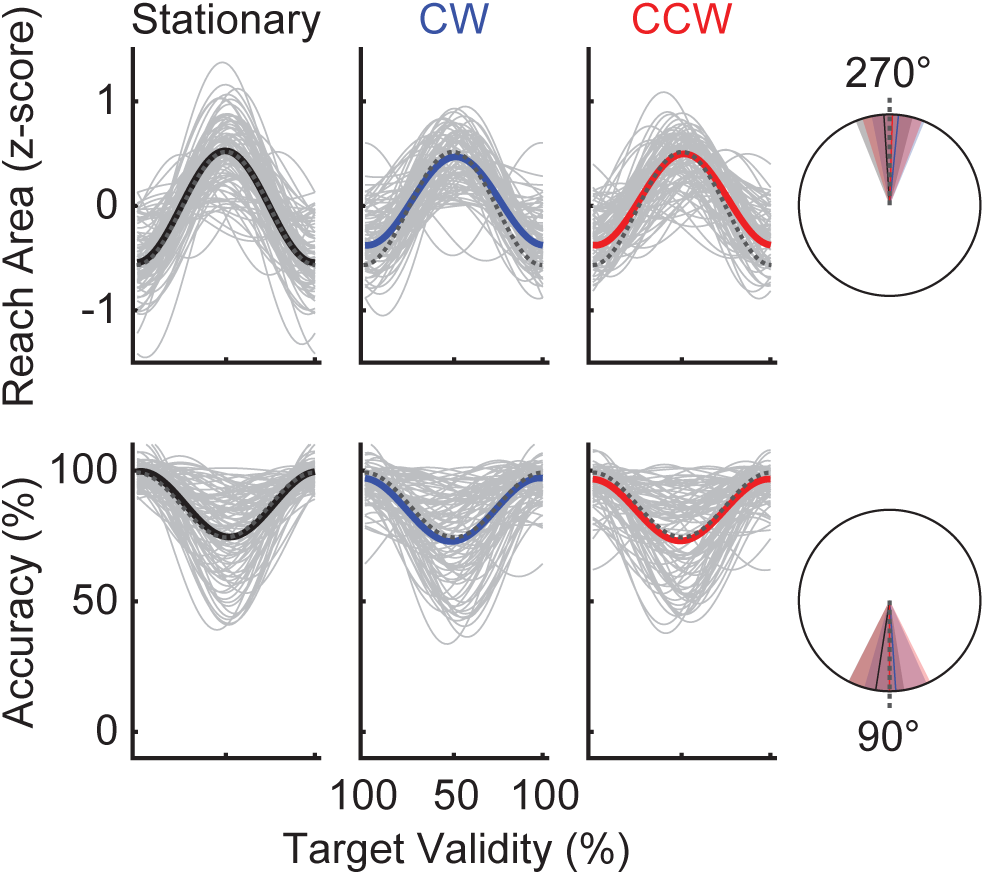
Sine wave models fit to reach area (top) and accuracy (bottom) data over target validity positions. As in Fig. 5, sine waves from each individual condition are in light grey, while a sine wave with average mean, amplitude, and phase parameters are in solid colors (black for stationary, blue for clockwise, red for counterclockwise). The average phase in each rotation condition and Bonferroni-corrected 95% confidence intervals are plotted along a circle, with the expected phase as a dashed line.

### 3.2. Additional main effects

Beyond the theoretically-motivated exploration of Phase parameters, we also examined differences in Mean and Amplitude for our sinusoidal parameter fits using a 3-factor repeated-measures ANOVA applied to each of reaction time, reach area, and accuracy. After correcting for the number of statistical tests (52), we found no significant effects of Experiment, nor any significant interactions in these data. Five main effects passed the adjusted significance threshold and are described below.

Analyses showed a main effect of rotation for the mean parameters estimated from reaction time data, *F*(1.21, 47.11) = 54.86, *p* = 5.8e-9. Bonferroni-corrected post-hoc comparisons showed mean parameters were lower in the stationary condition relative to the clockwise or counterclockwise conditions (*p*s < 8.00e-12), and that the clockwise and counter-clockwise conditions did not differ (*p* = 1.00). In other words, participants were faster to start moving when the stimulus was stationary relative to when it was moving.

Analyses also showed a main effect of rotation for the amplitude parameters estimated from normalized reach area data, *F*(1.61, 67.45) = 9.98, *p* = 4.39e-4. Bonferroni-corrected post-hoc comparisons showed amplitude parameters were higher in the stationary condition relative to the clockwise (*p* = 0.0003) or counterclockwise conditions (*p* = 0.002), and that the clockwise and counterclockwise conditions did not differ (*p* = 1.00). In other words, the difference between straight, confident reaches and indirect, conflicted reaches was larger for the stationary trials than the moving trials. This likely reflects that stationary trials’ probabilities were more discernible than rotating trials. Analyses revealed a main effect of target arrangement on the mean parameters estimated from accuracy data, *F*(1, 42) = 86.93, *p* = 8.69e-12. Post-hoc comparisons showed mean parameters were higher for horizontal targets relative to vertical targets (*p* = 4.83e-12), suggesting participants found horizontal trials easier than vertical trials.

Finally, analyses revealed a main effect of target arrangement for the amplitude parameters estimated from normalized reach area, *F*(1, 42) = 18.94, *p* = 8.46e-5, and accuracy data, *F*(1, 42) = 12.39, *p* = 0.001. Post-hoc comparisons showed amplitude parameters were higher for horizontal targets relative to vertical targets for reach area data (*p* = 8.47e-5). These results indicate that the change from straight to indirect reaches was larger for horizontal trials, likely because the hand started between the two targets for horizontal trials, but below the two targets for vertical trials. Conversely, amplitude parameters were higher for vertical targets relative to horizontal targets for accuracy data (*p* = 0.001). In other words, participants found horizontal trials easier than vertical trials. Essentially, accuracy was near 100% when probability was high for both horizontal and vertical trials, but vertical trials’ accuracy was much lower when probabilities neared 50%. This means that vertical trials have a larger amplitude to account for the decrease at 50% probability and subsequently have a lower mean.

## 4. Discussion

Here we assessed how well participants can plan an upcoming movement based on a dynamic, predictable stimulus that is not the target of action. A stationary or rotating stimulus determined the probability that each of two potential targets would be the ultimate target of a rapid reach-to-touch movement. Further, we used two different stimuli (box and arrow) to investigate processing differences in exogenous and endogenous attention systems. We questioned the extent to which the sensorimotor system integrates predictions about the future state of dynamic probabilistic information by examining movement trajectories.

We tested whether the sinusoidal pattern of reach area and accuracy was shifted in time relative to the rotation of the stimulus that determined target probability. We tested between three possible patterns of results (Fig. 3). According to the “delayed” prediction, transmission delays in the central nervous system would mean that reach area and accuracy would reflect the target probability at some time in the past. According to the “catch-up” prediction, information about target probability would be similarly delayed, but could still be used to correct online reach trajectories toward the final target more often. This catch-up prediction would appear as a delayed offset in the sinusoidal pattern of reach area relative to the probability-stimulus, but with less temporal offset for the sinusoidal pattern of accuracy. Finally, according to a “complete” prediction, participants would be able to successfully predict the future location of the probability-stimulus while accounting for their own reaction time, ultimately producing a sinusoidal pattern for both reach area and accuracy in lock-step with information about the final target probabilities. Our results support the notion of “complete” prediction (Fig. 3), wherein there is no temporal offset for patterns of reach area and accuracy between stationary, clockwise, and counterclockwise conditions. Overall, we show that, despite sensory and motor delays in the central nervous system, movement trajectories reflect target probability determined at movement onset. This was true for both the box and arrow experiment, suggesting that the prediction of probability from a non-target stimulus is not subject to changes due to a central versus more peripheral focus. This suggests that humans rapidly and accurately integrate visuospatial predictions from various non-target stimuli and can estimate their own reaction times to effectively guide action.

It has long been argued that one of the major roles of the brain is to produce movement (35, 50, 53, 54) and that this capacity, among others, involves prediction (55, 56). In short, several theories posit that the brain, rather than using the accumulation of bottom-up sensory cues to build a model of the world, instead builds predictions about the current state of the world and compares these predictions to incoming sensory information. The difference between the predicted sensory input and the actual sensory input—termed the “prediction error”—is used to continually update internal models of the world (55). Evidence for such predictive coding has been found for low-level sensory input (57, 58), as well as higher order cognitive functions (59, 60).

In addition to perception and cognition, the fundamental capacity for prediction is required for effective motor control, where appropriate motor commands are computed through the use of internal forward models (54). Forward models are a theoretical construct that can be used to predict, given a particular motor command, the sensory consequences of executing the action. Such prediction allows the brain to account for transmission and computational delays in the central and peripheral nervous systems, effectively providing for robustness in both real-time control and perception. There is good behavioural and neural evidence that the brain contains such internal models (61–63). For example, with respect to perception, we are unable to tickle ourselves because forward models can be used to inhibit sensations arising from self-motion (64). Likewise, with respect to control, the prediction of the sensory consequences of action can allow the brain to rapidly detect performance errors, and rapidly launch effective corrective actions as needed. A forward model is useful especially when generating interceptive actions. How we use an internal prediction model for interceptive actions was tested by Soechting, Juveli, & Rao (2009) using a model that explained finger movements during interception of a randomly moving target on a screen (65). They found that the finger’s position within 100 ms of movement onset reflected anticipatory predictions in advance of the target’s location, similar to our reach area results. However, Soechting *et al*. (2009) conclude that only “directly observable quantities” like target position and velocity are integrated into an internal prediction model, while higher order properties like statistical features of motion (i.e., sinusoidal motion laws) are not dynamically refined. In contrast, our results suggest that some *unobservable* quantities, in this case target probability derived from a rotating stimulus, do indeed directly impact real-time predictions.

In this study, we used movement trajectories to reveal the sensitivity to changes in target probability. Previous work has shown that trajectories are thought to be a real-time readout of several cognitive variables, shown in behaviours such as changes of mind (4, 5), or moment-to-moment fluctuations throughout movement (24, 66). Here, we show that curved reach trajectories (i.e., those with large reach areas) reflect uncertainty about the predicted target position, while relatively straighter reach trajectories (i.e., those with smaller reach areas) reflect more certainty about target predictions. We provide Figure 7 as a useful descriptive tool demonstrating the effect of the position of our dynamic probability stimulus at the time of the go-signal on average participant trajectories. For stationary stimuli, reach trajectories are most curved when the stimulus is positioned half-way between the two targets (50% target probability, Fig. 7a, middle panels of top and bottom rows), whereas for rotating stimuli reach trajectories are most curved when the stimulus is moving *toward* 50% probability at the go-cue (top left and bottom right corners for Fig. 7b, CW; top right and bottom left corners for Fig. 7c, CCW). Overall, this suggests that participants are successfully predicting the future probability of both potential targets, and planning their movements accordingly.

**Fig. 7.**
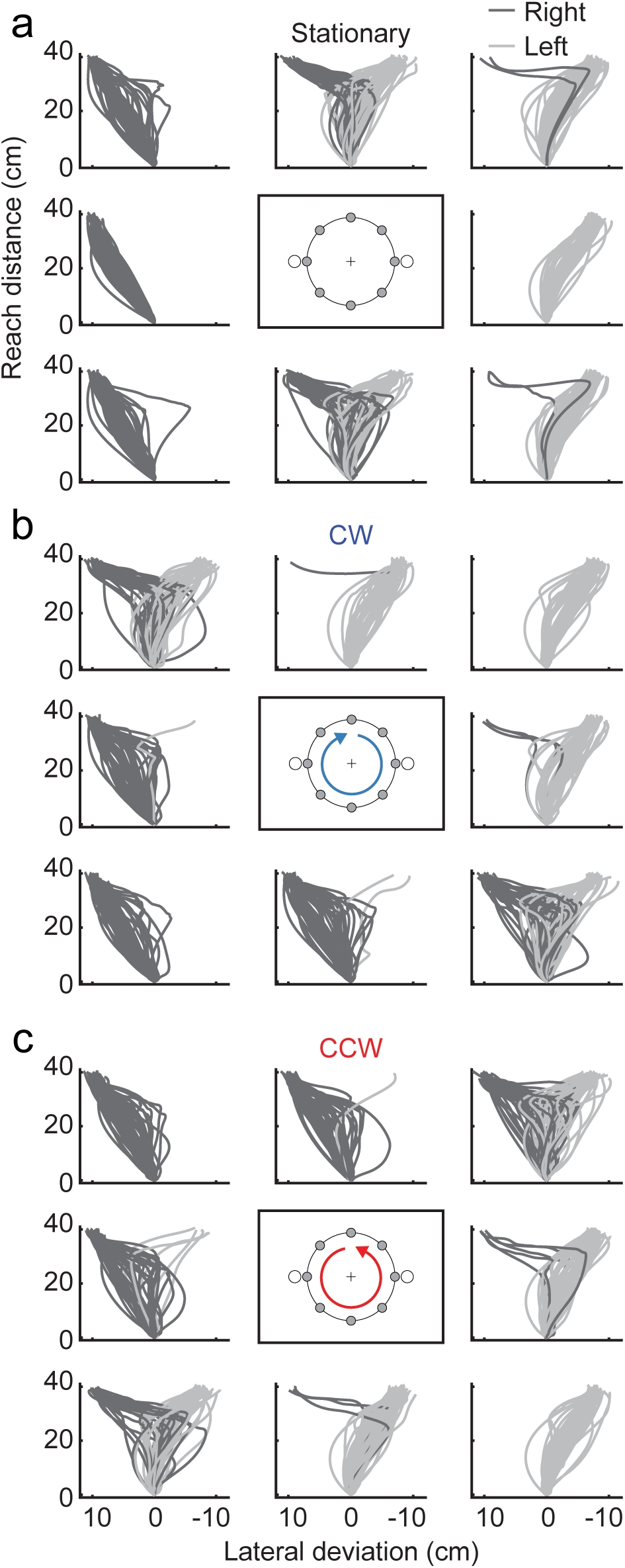
Average correct reach trajectories for each participant when reaches ended left (dark grey) and right (light grey) on trials where targets were arranged horizontally. Reaches are plotted in one of the eight trigger positions along the circle corresponding to the probability-stimulus location when the go-signal was presented. These data are intended to be qualitative descriptors of the overall reaching patterns by condition. For (a) Stationary trials, reaches tended to be straightest when the probability-stimulus is at the left or rightmost trigger position at the go-signal, and more curved when at the top and bottommost trigger position. For (b) Clockwise and (c) Counterclockwise trials, this pattern is shifted indicating that participants were anticipating the future location of the probability-stimulus.

Behavioural measures such as reaction time, accuracy, and movement trajectories are often thought to index the same internal cognitive processes (31). However, these behaviors are measured at different times. For instance, reaction time may reveal cognitive variables several hundred milliseconds before accuracy, particularly when a reaching movement separates the two. Differences between these measures may reveal the evolution of cognition over the course of a trial, especially in dynamic environments.

Here we see a dissociation between reaction time and measures of reach area and accuracy. The pattern of reaction time on clockwise and counterclockwise trials is out of phase with the pattern of reaction time on stationary trials—results not observed for reach area and accuracy. On one hand, this difference could reflect that reaction time is indexed at an earlier point in the trial than accuracy and most of the movement trajectory. This might suggest that the internal prediction of target probability is still evolving when reaction time is measured, while predictions of target probability are accurate at the time movement and accuracy are measured. Some models explicitly theorize that reaction time, movement trajectories, and accuracy can be explained in some tasks from the same internal decision variable (4). However, these measures, while similar in some tasks, may arise from distinct computation. Such differences may also explain the discrepancy between reaction time, reach area, and accuracy in the current results. Finally, it is also possible participants are adjusting reaction times to improve visuomotor accuracy in trials with greater uncertainty (43). When trials are uncertain, longer reaction times may be used to accumulate more sensory evidence to guide their decision. Overall, however, these results suggest more work needs to be done to determine if reaction time and movement variables in a reach decision task reflect common or separate cognitive processes.

In the present study, we demonstrated that humans are able to accurately predict future states from a predictable, dynamic, non-target object and account for sensorimotor delays to guide rapid reaching movements. Such predictions are likely a key part of neural computation within and between different systems of the brain. The results of this study speak to one key part of how humans are able to carry out actions in complex and dynamic environments.

## Data and code availability

Videos of the task, data, and analysis code are publicly available at the following website: https://osf.io/rt5xv/.

## ACKNOWLEDGEMENTS

This research was supported by the National Science and Engineering Research Council of Canada (NSERC) and the Killam Trusts. This preprint article is formatted using a LATEX template by Ricardo Henriques.

## AUTHOR CONTRIBUTIONS

J.P.G. and C.S.C. conceptualized the experiments and collected data. All authors analyzed the data and were involved in writing, reviewing, and editing.

